# Comparative genomic and pan-genomic characteristion of *Staphylococcus epidermidis* from different sources unveils the molecular basis and potential biomarkers of pathogenic strains

**DOI:** 10.1101/2021.08.12.456181

**Authors:** Shudan Lin, Bianjin Sun, Xinrui Shi, Yi Xu, Yunfeng Gu, Xiaobin Gu, Xueli Ma, Tian Wan, Jie Xu, Jianzhong Su, Yongliang Lou, Meiqin Zheng

## Abstract

Coagulase-negative *Staphylococcus* (CoNS) is the most common pathogen causing traumatic endophthalmitis, *Staphylococcus epidermidis* is the most common species which colonizes human skin, eye surfaces and nasal cavity and is the main cause of nosocomial infection, specially foreign body-related bloodstream infections (FBR-BSIs). Although some studies have reported the genome characteristics of *S. epidermidis*, a comprehensive understanding of its pathogenicity and the genome of ocular trauma-sourced strains is still lacking. In this study, we sequenced, analyzed and reported the whole genomes of 11 ocular trauma-sourced samples of *S. epidermidis* that caused traumatic endophthalmitis. By integrating publicly available genomes, we obtained a total of 187 *S. epidermidis* samples from healthy and diseased eyes, skin, respiratory tract and blood. Combined with pangenome, phylogenetic and comparative genomic analyses, our study supported that *S. epidermidis*, regardless of niche source, exhibits two founder lineages with different pathogenicities. Moreover, we identified potential biomarkers associated with the virulence of *S. epidermidis*, namely, *essD*, *uhpt*, *sdrF*, *sdrG*, *fbe* and *icaABCDR*. The *essD* and *uhpt* genes have high homology with *esaD* and *hpt* in *Staphylococcus aureus*, showing that the genomes of *S. epidermidis* and *S. aureus* may have communicated during evolution, while the *sdrF*, *sdrG*, *fbe*, and *icaABCDR* genes are related to biofilm formation. Compared to *S. epidermidis* from blood sources, ocular-sourced strains causing intraocular infection had no direct relationship with biofilm formation. In conclusion, this study not only provided additional data resources for studies on *S. epidermidis*is, but also improved understanding of the evolution and pathogenicity of different source strains.

**Importants:** In this study, we comprehensively analysied and reported the whole genome sequence (WGS) of the strains that caused traumatic endophthalmitis. Through comparative genomic analyses among 187 *S. epidermidis* samples from healthy and diseased eyes, skin, respiratory tract and blood, we identified 10 potential biomarkers of pathogenic strains. Finally, we revealed *S. epidermidis*-induced traumatic endophthalmitis may be independent of biofilm formation. Overall, our data may facilitate comparative research of *S. epidermidis* and provide clinical value for identifying pathogenic or contaminating strains.

## Introduction

Coagulase-negative *staphylococci* (CoNS) usually live on human skin, can be cultivated from every individual, even healthy persons, and represent one of the major nosocomial pathogens(1). As recorded within the U.S. nationwide Surveillance and Control of Pathogens of Epidemiological Importance (SCOPE) database, the most common isolate recovered from nosocomial bloodstream infections was CoNS (31%) at the end of a 7-year period(2). Additionally, CoNS preferentially colonized the nasal cavity(97%)(3) and ocular surface (60%)(4)(5). *Staphylococcus epidermidis* is the most common CoNS(6), and it plays a central role in the skin microbiota; for example, it can protect against colonization by skin pathogens(7)(8), maintain the ecological balance of human skin flora(9) and modulate the immune system(10)(11). Once it breaches the skin surface and enters the bloodstream, however, it is considered pathogenic. As the second most common cause of nosocomial infections(12), *S. epidermidis* not only accounts substantially for foreign body-related infections(13) but also causes many eye infections(14)(15)(16)(17), such as bacterial endophthalmitis, keratitis, conjunctivitis and blepharitis, mainly because of ocular trauma(17), ocular surgery, ocular inflammation and the use of contact lenses(14)(17). The data obtained in different ways all showed that CoNS, including *S. epidermidis* (more than 30%), was the most common pathogen causing traumatic endophthalmitis, and more than 50% of patients could not recover effective visual acuity (0.1), among which more than 20% completely lost visual acuity. The different roles of *S. epidermidis* in health and disease give it an important and central role in the human microbiota. Although a high positive rate of *S. epidermidis* was detected in clinical specimens, whether they represent true infection or only colonization/contamination remains to be discussed.

Although much less is known regarding the potential of *S. epidermidis* to cause outbreaks, the advantage of CoNS isolates showed (multiple) resistance to antibiotics (18)(19)(20), as well as their ability to produce biofilms(21)(22), strongly suggesting that modern medicine facilitated the selection process, mainly by the (over)use of antibiotics and the insertion of foreign body devices(13). The biofilm formation and antibiotic resistance of *S. epidermidis* contributed to the occurrence and persistence of clinical infections(23). Studies conducted worldwide have shown that in clinical specimens, especially in medical device specimens, the isolation rate of *S. epidermidis* is very high(24)(25)(26). *S. epidermidis* forms biofilms on medical devices, such as contact lenses, catheters, and artificial heart valves(27). The detachment of bacterial cells from the biofilm on these devices can lead to bacteremia, increasing morbidity and potential mortality(27). The biofilm formation of *S. epidermidis* was caused by initial attachment by surface proteins (staphylococcal surface proteins *Ssp-1*, *Ssp-2*, Bap homolog protein *Bhp*, autolysin E *AtlE*) and indirect binding, which was accomplished by identifying the microbial surface components of adhesion matrix molecules (MSCRAMMs), such as fibrinogen-binding protein (*SdrG*, *Fbe*) or polysaccharide intracellular adhesion (*ica* Luci)(28). It is well known that biofilms are resistant to antibacterial drugs(27).

Antibiotic resistance significantly complicates treatment and increases medical costs(29)(30). Mobile genetic elements, such as multidrug resistance binding plasmids and the set of recombinase genes (*ccr*) called staphylococcal chromosome cassette *mec* (SCC*mec*)(31), enable frequent transfer of β-lactam resistance and adaptation to antibiotic selection pressure by *S. epidermidis*(32)(33). The gene *mecA* is present on SCC*mec* and encodes the penicillin-binding protein PBP2a(34). The alteration in PBP2a may confer resistance to methicillin(35). Although methicillin is not used to treat eye infections, it is known that increasing resistance to methicillin can significantly promote the spread and persistence of multidrug-resistant strains in specific environments (36)(37)(38).

Recently, with the development of high-throughput sequencing technology, the genome of *S. epidermidis* has also been extensively studied. Conlan et al first analyzed the full pangenome of *S. epidermidis*, including both commensal and nosocomial isolates(27). Sharma et al reported the presence of diverse lineages of *S. epidermidis* isolates in healthy individuals from two geographically diverse locations in India and North America(39). Su et al selected isolates from human and environmental sources from databases to identify genomic determinants associated with the diversity and of *S. epidermidis* strains and their adaptation to various environments(40). Zhou et al characterized the evolutionary trajectory and functional distribution of *S. epidermidis*(41). Microorganisms can be shaped by different host-specific factors, such as disease and health status, however, because *S. epidermidis* is an opportunistic pathogen with a high isolate rate, we remained unsure whether the detected strains were true infection or only contamination, especially in the case of respiratory tract samples, which were generally considered to be contaminated and were not reported. Thus, the pathogenic condition of *S. epidermidis* from different host health statuses should be fully understood. Moreover, although Kirstahler et al reported six whole genomes of *S. epidermidis* from vitreous humor(42), a comprehensive understanding of the genome of ocular strains, particularly ocular trauma-sourced strains, was still lacking, and genetic differences in strain sources from different host niches remained unclear.

In this study, we sequenced the whole genomes of 11 ocular trauma-sourced *S. epidermidis* isolates. Through incorporating publicly available genomes, we finally obtained 187 *S. epidermidis* isolates from eyes, skin, respiratory tract and blood of healthy and diseased hosts. Through whole-genome sequence analysis and pangenome analysis, we performed a detailed analysis of the host health status diversity and within-individual sources of *S. epidermidis*, and we comprehensively analyzed and reported the whole-genome sequence (WGS) of *S. epidermidis* that causes traumatic endophthalmitis. We used phylogenetic analysis and comparative genomics to reveal the evolutionary relationships of different sources, exploring how different host niches may shape the genetic diversity of *S. epidermidis*. Our study revealed a marked association between evolutionary lineage and host health states. Regardless of niche source, *S. epidermidis* showed two founder lineages that had different pathogenicity. We also identified biomarkers related to *S. epidermidis* pathogenicity. Compared to blood-sourced *S. epidermidis*, traumatic endophthalmitis strains carried different virulence genes, and causing intraocular infection may be independent of biofilm formation. Overall, our study revealed the genetic diversity and pathogenicity of *S. epidermidis* and accomplished a comprehensive comparative analysis of different source genomes.

## Materials and methods

### Strains

This study analyzed the whole genomes of 187 *Staphylococcus epidermidis* isolates, including 11 ocular strain isolates from the Department of Laboratories, Eye Hospital of Wenzhou Medical University, Wenzhou, China, and 176 available genome sequences downloaded from the GenBank database(43) of the National Center for Biotechnology Information (NCBI) (http://www.ncbi.nlm.nih.gov). The 187 *S. epidermidis* strains were selected to represent known diversity within isolation sources representing human and host health states. For strain sources, we collected 17 ocular *S. epidermidis* strains, 46 blood-sourced strains, 22 respiratory strains, 46 *S. epidermidis* skin-sourced strains, and 46 clinical strains with unknown sources of host niches classified as group clinics. of the remaining *S. epidermidis* strains belonged to the “Others” group. The “Respiratory” group consists of isolates from the nares, lungs, pharyngeal exudate, sputum and bronchoalveolar lavage. Catheter and oral isolates and the reference genomes of strains ATCC14990 and RP62A were classified in the “Others” group. A total of 19 and 75 strains had a definite health or disease state, respectively.

### Whole-Genome Sequencing, Assembly, Gene Predictions and Functional Annotations

The 11 *S. epidermidis* isolates collected from ocular sources were cultured in 5 ml of brain heart infusion broth (BHI) +5% fetal bovine serum (FBS, Gibco) and shaken at 250 rpm and 37°C for 12–16 h. DNA was extracted using a GENEray Bacterial Genome DNA Mini Kit (GENEray, Shanghai, China) following the manufacturer’s protocol. Whole-genome sequencing was performed by Berry Genomics Co., Ltd, Beijing, China using PacBio SMRT Technology. SMRTbell libraries were prepared using the SMRTbell Express Template Prep Kit 2.0 (PacBio kit). Long read data were assembled by canu 2.1.1(44), and assembly polisher was used with pbmm2 (v1.4.0) and gcpp (v2.0.0) via the bioconda package. Quast (v5.0.2)(45), checkM (v1.1.3)(46) and busco (v5.0.0)(47) were used to assess the quality of assembled genomes. The genes of all genomes, including the 11 ocular *S. epidermidis* genomes that we sequenced and 176 obtained from NCBI, were predicted and annotated by Prokka (v1.13)(48).

### Pan-Genome Analysis

Pangenome analysis was carried out for all 187 genomes and isolate genomes from different niches by BPGA v1.3(49) using default parameters and 50% identity as the cutoff. The annotation files generated by Prokka were provided to BPGA as an input. The COGs of the core sequence, accessory sequence and unique sequence after Pangenome analysis were further verified by performing functional annotations in the EggNOG Database (v5.0)(50) via eggnog-mapper (v2.0)(51) with the Diamond parameter.

### Characterization of Strains

kSNP3.1(52) was used to build the phylogenetic tree, a validated method without alignment by 19 k-mer length, and a total of 8760 core SNPs were identified. Then, a maximum-likelihood core SNP tree was constructed by RAxML (v8.2.12)(53) with 100 bootstraps. The phylogenetic tree was visualized and beautified by online itol (v6.0) (https://itol.embl.de/)(54).

Multilocus sequence typing (MLST) of 187 *S. epidermidis* strains was performed with the MLST 2.0 online server (https://cge.cbs.dtu.dk/services/MLST/)(55). Online SCC*mec*Finder 1.2 (https://cge.cbs.dtu.dk/services/SCCmecFinder/) was used to identify SCC*mec* elements in sequenced *S. epidermidis* isolates.

ResFinder(56) and Resistance Gene Identifier (RGI) of the Comprehensive Antibiotic Resistance Database (CARD)(57) were employed to identify antimicrobial resistance genes (AMRs). The resistance genes were verified by the paper diffusion method (MHA) and broth dilution method. The BLASTp program was used to search all protein sequences of 187 *S. epidermidis* strains against the Virulence Factor Database (VFDB) (http://www.mgc.ac.cn/VFs/)(58). Compared with the virulence genes in the database at an e-value < 1e-10, only query genes with an identity higher than 40% and a coverage higher than 70% were considered potential virulence genes(59). Virulence factor functional annotations were based on the categories and subcategories presented in VFDB. An alignment for each of the extracted candidate virulence determinant genes was constructed using Clustal Omega(60) and visualized by ESPript(61).

### Statistical Analyses

The significance of core gene and accessory gene abundance in COG categories was examined using Fisher’s exact test. The disease-associated and ocular-associated predicted accessory genes with known function and annotated virulence genes were analyzed using Fisher’s exact tests and the FDR correction of P values. All statistical analyses were carried out using the R package (version: 4.0.2). A P-value of < 0.05 was regarded as statistically significant.

### Data availability

The sample and sequence data obtained in this study have been submitted to the NCBI BioSample and Sequence Read Archive (SRA) under BioProject accession number PRJNA753005.

## Results

### General Features of Sequenced Ocular Isolate Genomes

Eleven *Staphylococcus epidermidis* isolates from *S. epidermidis*-infected ocular endophthalmitis were sequenced and assembled. We used pbmm2 to map the draft genomes to raw subreads and ultimately counted the coverage depth. High-quality data were obtained with a mean coverage depth >5000X and number of contigs in the range of 1–6. Annotation of these isolates revealed predicted CDSs (from coding sequences) ranging from 2,247 to 2,692. The sequencing and annotation details of the 11 ocular isolates are listed in **Table 1**. The 11 draft genomes were 2.58±0.1 Mb in size and 32.14% GC on average. It seemed that the features of sequenced ocular isolate genomes were not significantly different from those obtained from skin and blood. We chose a representative assembled ocular strain genome and drew a genomics map with Circos(62) to illustrate the genome characterization (**Figure 1**).

**Table1.** Genomic characteristics of 11 ocular trauma-source *Staphylococcus epidermidis*

|  | Mapping<br>Reads | Unmappin<br>-g Reads | Mean<br>Coverage<br>Depth | Coverage<br>Depth<10<br>(%) | GC<br>content | Contig<br>number | Genome<br>size<br>(Mb) | CDS | tRNA | rRNA |
| --- | --- | --- | --- | --- | --- | --- | --- | --- | --- | --- |
| O106 | 1063834 | 0 | 4524 | <0.001% | 32.27% | 2 | 2.54 | 2360 | 60 | 19 |
| O107 | 1635910 | 0 | 6830 | 0 | 32.15% | 3 | 2.56 | 2355 | 60 | 19 |
| O108 | 1148008 | 0 | 5347 | 0 | 32.24% | 1 | 2.49 | 2247 | 60 | 22 |
| O109 | 1457938 | 0 | 6404 | 0.0023% | 32.10% | 3 | 2.56 | 2391 | 59 | 19 |
| O110 | 1523429 | 0 | 6511 | 0 | 32.05% | 3 | 2.67 | 2446 | 60 | 19 |
| O112 | 1629802 | 0 | 6936 | 0 | 32.17% | 2 | 2.54 | 2313 | 59 | 19 |
| O113 | 1495273 | 0 | 6420 | 0.016% | 32.04% | 4 | 2.62 | 2455 | 60 | 19 |
| O114 | 1485509 | 0 | 6590 | <0.001% | 32.15% | 4 | 2.58 | 2345 | 60 | 19 |
| O115 | 988780 | 0 | 4438 | 0 | 32.15% | 2 | 2.51 | 2272 | 60 | 19 |
| O116 | 1110019 | 0 | 4641 | 0.1% | 32.01% | 6 | 2.79 | 2692 | 60 | 19 |
| O117 | 1130833 | 0 | 5414 | <0.001 | 32.27% | 1 | 2.48 | 2265 | 61 | 19 |

**FIGURE 1.**
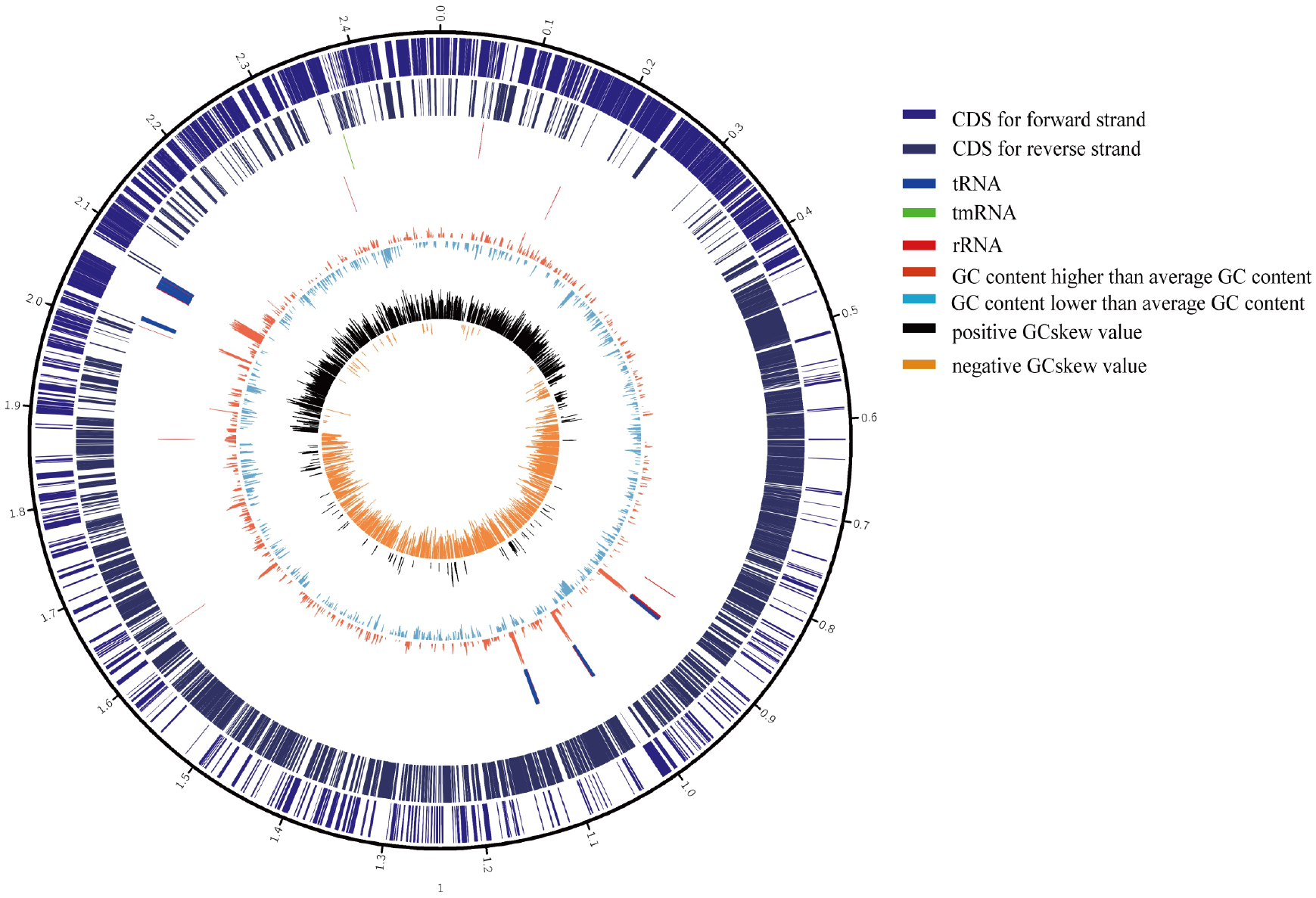
Circular genome maps of representative ocular strain. From the outer to the inner circle: (1) scale marks of genomes; (2) assigned COG classes of protein-coding genes (CDSs) on the forward strand; (3) reverse strand CDSs; (4) tRNA (blue) and rRNA (red) genes on the forward strand; (5) tRNA (blue) and rRNA (red) genes on the reversed strand; (6) GC content (swell fire red/sky blue indicates higher/lower G + C compared with the average G + C content); (7) GC skew (black/orange indicate positive/negative values).

### Pan-Genome and Functional Characterization of *S. epidermidis* from Different Sources

Although many studies have shown that pangenomes characterize *S. epidermidis*(27), core and pangenome features of ocular strains are lacking. Thus, to clarify the pangenomic characteristics of *S. epidermidis* from ocular sources and discrepancies among different sources, we determined general genetic similarities and differences within ocular *S. epidermidis* and all 187 strains. Gene accumulation curves(63) **(Figure 2A and Figure 2B)** showed that the number of core genomes fits an exponential decay curve that plateaus at 749 and 1,931 genes, respectively, while the pangenome data fit a power law curve (y=a·x^b^), indicating an open pangenome where each genome sequence added a number of new genes, as reported(27). To a certain degree, the capacity of acquiring exogenous DNA of the organism partially determines the pangenome state (“open” or “close”)(64), especially for species living in bacterial communities, such as skin inhabitants.

**FIGURE 2.**
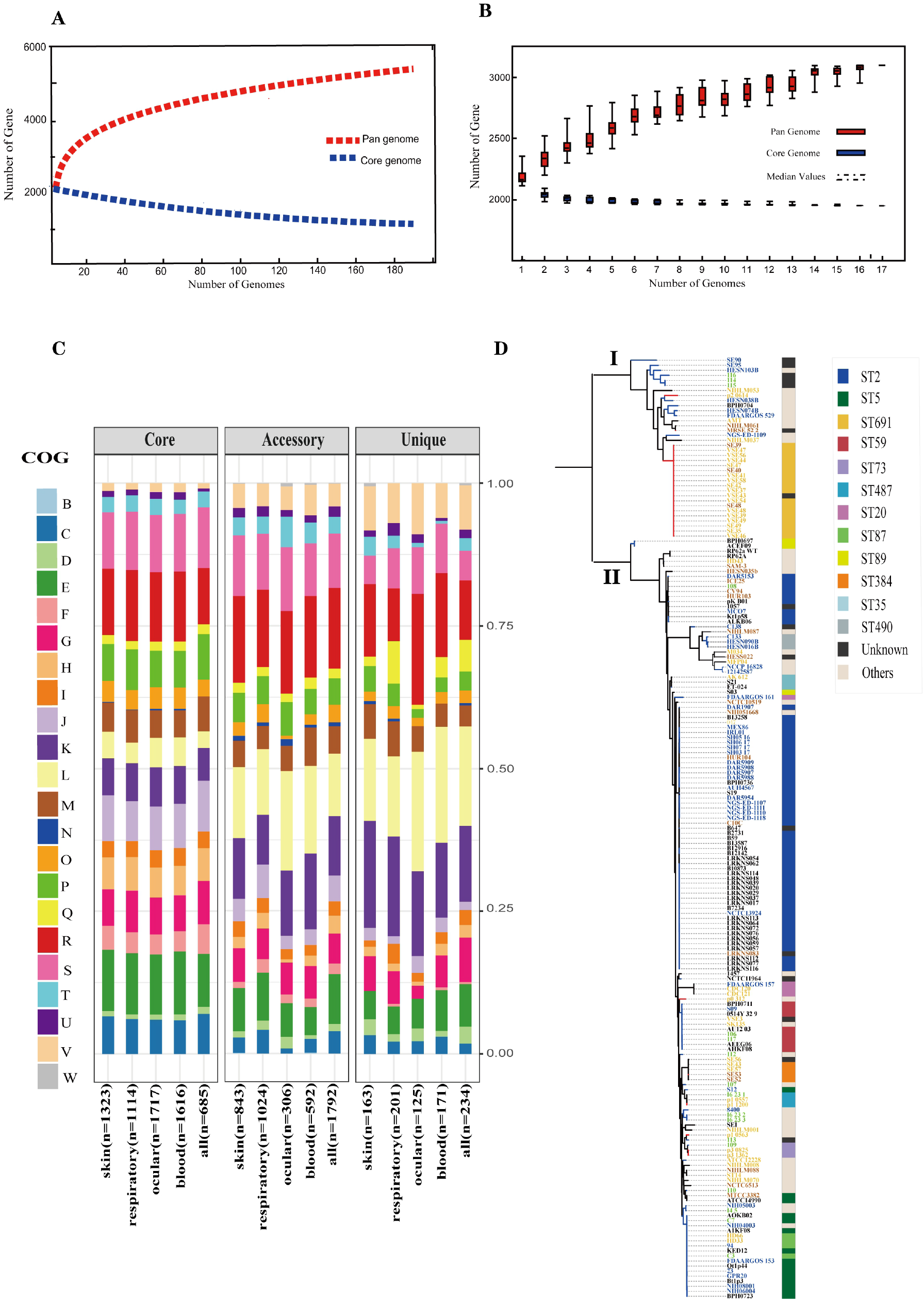
Pan-genome and phylogenetic feature of S. epidermidis from different host niches and health state. A.Gene accumulation curves for pangenome (red) and core-genome (blue) of collected all strains. B. Gene accumulation curves for pangenome (red) and core-genome (blue) of ocular sources strains. C. COG functional categories from the pan-genomes witihin strains from different niches. Involved COG categories are as follows:[B]Chromatin structure & dynamics; [C]Energy production & conversion; [D]Cell cycle control, cell division, chromosome partitioning; [E]Amino acid transport & metabolism; [F]Nucleotide transport & metabolism; [G]Carbohydrate transport & metabolism; [H]Coenzyme transport & metabolism; [I]Lipid transport & metabolism; [J]Translation, ribosomal structure & biogenesis; [K]Transcription; [L]Replication, recombination & repair; [M]Cell wall/membrane/envelope biogenesis; [N]Cell motility; [O]Post−translational modification, protein turnover & chaperones; [P]Inorganic ion transport & metabolism;[Q]Secondary metabolites biosynthesis, transport & catabolismand; [T]Signal transduction mechanisms; [U]Intracellular trafficking, secretion & vesicular transport; [V]Defense mechanisms; [W]Extracellular structures. Poorly characterized COG categories contains [R]General function prediction only and [S]Function unknown. D. Phylogenetic core SNP maximum likelihood tree was constructed for 187 genomes. The blue branch represent strains from diseased hosts while heathy source strains put color on red. Different color of strains ID stand for different host niches: blood (blue), ocular (green), skins (yellow) and respiratory tract (brick-red).Legends on the left stand for colors of sequence type(ST) from multi-locus sequence typing.

**FIGURE 3.**
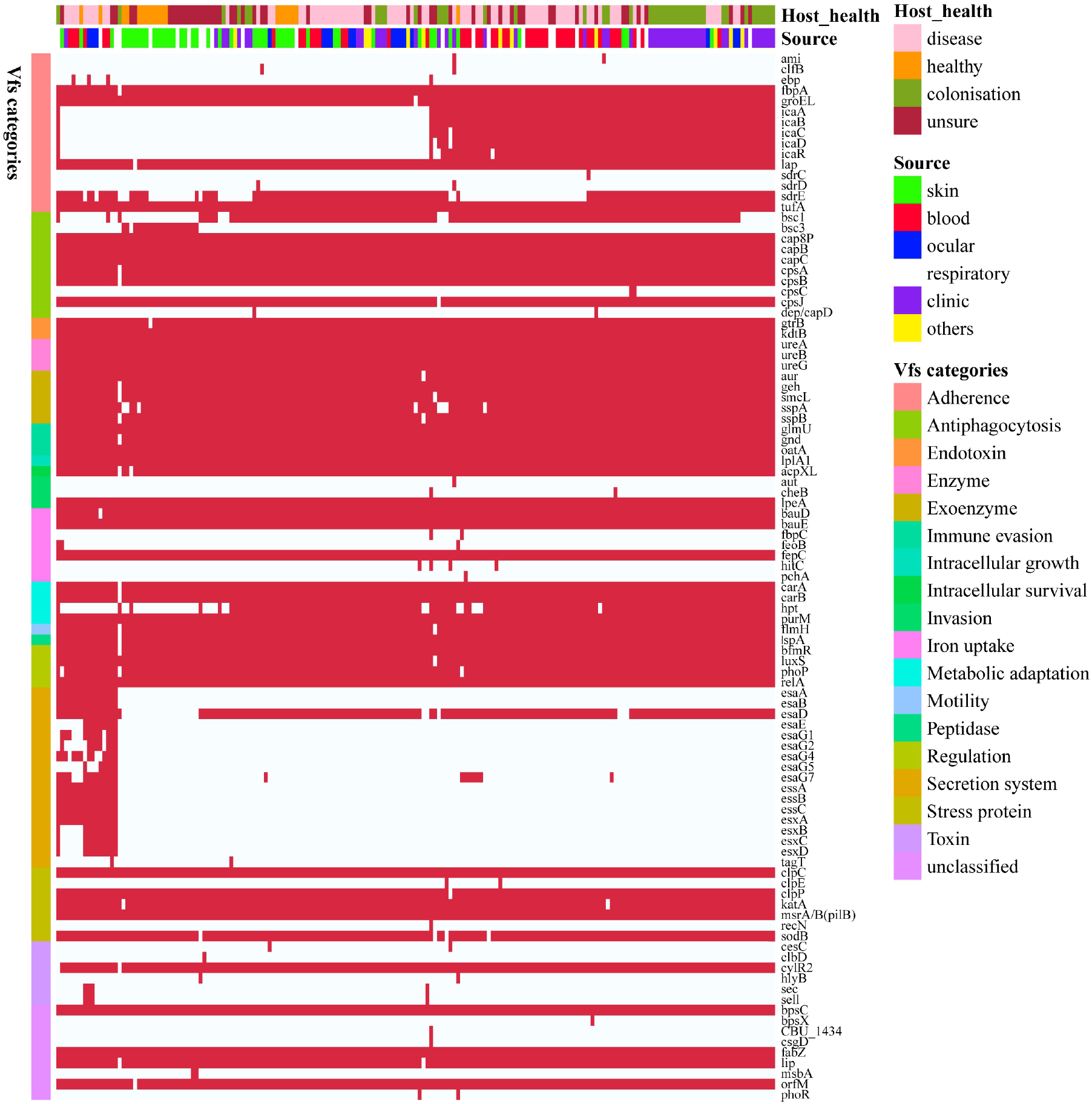
Heatmap of virulence genes in all 187 S.epidermidis strains. The red square indicates the presence of the gene, while the white square indicates absence. Legends on the right stand for colors of different host healthy state, isolates source and virulence factor categories.

The function of the genes within the pangenome of whole collected and different source strains was investigated by assigning all gene clusters to clusters of orthologous groups (COGs)(65) **(Figure 2C)**. For entirety, the results revealed that there were a total of 685/749 core genes, 1,792/2,783 accessory genes and 234/468 unique genes among all 187 *S. epidermidis* strains annotated to COG categories. Specifically, by Fisher’s exact test (FDR < 0.05), 5 of 22 COG categories were significantly enriched in core genes and almost associated with metabolism and biogenesis: energy production and conversion, nucleotide transport and metabolism, coenzyme transport and metabolism, translation, ribosomal structure and biogenesis and inorganic ion transport and metabolism.

Additionally, while replication, recombination and repair were enriched in both accessory and unique genes, secondary metabolite biosynthesis, transport and catabolism and defense mechanisms were enriched in unique genes. Second, we compared the enrichment of COG function among strains from different sources, including the blood, eyes, respiratory tract, and skin, and showed similar results: core genes related to metabolism and biogenesis. However, the difference from the above is that defense mechanisms were enriched in accessory genes of all strains except eye-sourced ones. In addition, transcription was significantly enriched in accessory genes of blood-sourced strains, while other sources were enriched in unique genes. In summary, compared to the core genes enriched in biogenesis and metabolism, the enrichment of replication, recombination and repair, defense mechanisms, and transcription among accessory genes and unique genes was driven by diversity in recombinase and integrase, ABC-type multidrug transporters, and transcriptional regulators, respectively. These abundant genes often appear to transfer horizontally between strains, leading to the spread of virulence and resistance genes between strains, which affects the pathogenicity of bacteria(66).

### Phylogenetic Relationship and Associated Typing among *S. epidermidis*

To infer the phylogenetic relationship of the *S. epidermidis* strains from different sources, we used 8760 core SNPs to build a single-nucleotide polymorphism-based phylogenetic tree. As expected, the 187 *S. epidermidis* isolates formed two distinct groups termed I and Ⅱ, as previously reported(27)(41) (**Figure 2D**), suggesting the presence of two founder lineages. Moreover, we found that ST2 and ST5 strains were present in Cluster Ⅱ and correlated with definite disease hosts or clinical strains, while ST691 strains were present in Cluster I and were related to healthy skin with the same genetic distance.

It is worth mentioning that although the ocular isolates we examined are varied in phylogeny, they all had a close distance to different sources from diseased hosts in both clades, which proved that the ocular strains we collected were pathogenic. However, the ST typing of ocular strains was highly diverse and included rare specimens: 4/11 strains we collected were unknown.

*S. epidermidis* contains staphylococcal chromosome cassette *mec* (SCC*mec*), called methicillin-resistant *S. epidermidis* (MRSE)(27). SCC*mec* is a mobile genetic element defined by combinations of *mec* gene complexes, cassette recombinases and accessory genes and carries the central determinant for broad-spectrum beta-lactam resistance encoded by the *mecA* gene, a mobile genetic element of *Staphylococcus* species(27)(67). We predicted the SCC*mec* element (SCC*mec* type) and found that it existed in 94/187 strains. Interestingly, almost all SCC*mec*-positive strains (96.8%, 91/94) were in Cluster II, indicating that compared to SCC*mec*-negative strains, *S. epidermidis* carrying the SCC*mec* element may be more pathogenic.

### The Virulence Characteristics of *S. epidermidis*

Although *S. epidermidis* harbors few classical virulence determinants(6), it is an opportunistic pathogen with many virulence factors. To analyze the different pathogenic potentials within *S. epidermidis* from different sources, 99 genetic loci were identified to be related to virulence based on the virulence factor database (VFDB), which was grouped into 18 categories **(Figure 4)**, including 21 (21/99, 21.2%) virulence genes that coexisted in all *S. epidermidis* genomes. Surprisingly, we identified several virulence genes, *icaABCDR*, *hpt* and *esaD,* that were significantly associated with *S. epidermidis* from diseased sources (Fisher’s exact test, FDR < 0.01). Polysaccharide intercellular adhesion (*icaABCD*) genes that encode biofilm-associated genes for poly-N-acetylglucosamine synthesis were found in 48% (36/75) of the diseased isolates; notably, only 1 of 19 healthy strains contained the *icaABCD* gene, which disagreed with previous studies that found this gene in 60% of commensal isolates. By pairwise comparison of the enrichment of virulence gene in ocular-sourced strains and strains from other niches, we found 30 *icaABCD* genes in 46 blood-sourced strains and 3 in 17 ocular-sourced strains. We also found the Ser-Asp-rich proteins *sdrE* to be enriched in ocular-sourced strains (16/17) compared to blood-sourced strains (24/46) (Fisher’ exact test, FDR < 0.05). Two toxin genes, sell and sec, which were present in only two strains of blood-sourced *S. epidermidis* (SE90 and SE95), as reported previously(68), were also identified in two ocular strains, combining the results of phylogenetics showing that the strains had a close evolutionary distance, implying that they may have the similar founder lineages.

**FIGURE 4.**
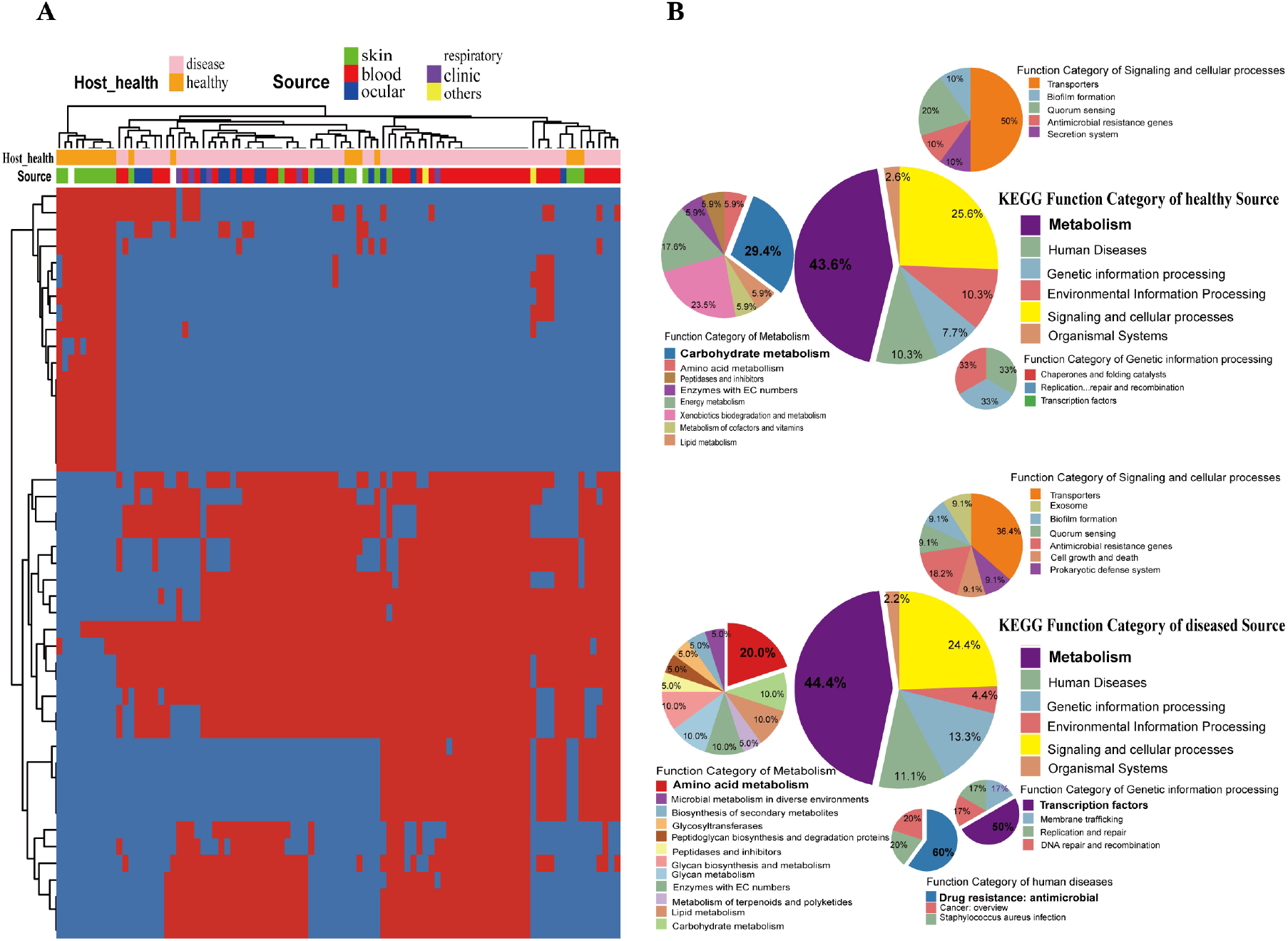
Gene function difference between healthy and disease source genomes. A. Heatmap of enriched genes (fisher’exct test, adjust P<0.05) between diseased and healthy sources strains. Only the accessory genes were shown. Gene clusters present in all genomes (core gene) or present in only a single genome (unique genes) are omitted. The red color stands for genes that existed and the blue color for missing ones. B. Pie graph of KEGG function related with enriched genes from diffrent health source. Bigger pie represented KEGG function proportion of 45 enriched genes showed in figure3A, while three smaller pie were account for function categories of metabiosim, genetic information process, human diseases, signaling and celluar processes, respectively.

### Gene Differences between Healthy and Disease Source Genomes

*Staphylococcus epidermidis* is a coagulase-negative and gram-positive *staphylococcus* that is part of the skin, mucosa microflora(69). They also had high relative abundances and high positivity rates on the ocular surface. It is the second leading cause of nosocomial infections(70). Within the 187 strain genomes we analyzed, 75 were definitely from diseased hosts and 19 were from healthy individuals. To investigate whether the presence of specific genes was significantly correlated with different sources of strains, including isolates from different healthy hosts. We devoted to accessory genes, which were defined to be gene clusters neither present in all genomes nor exit in only a single genome. Distinctly, the genes were divided into two clusters, which strictly differentiated the disease group and healthy isolates (Fisher’s exact test, FDR < 0.05) (**Figure 4A**). Ulteriorly, the two groups of genes were functionally classified by KEGG function (**Figure 4B**). Both congruently, metabolism accounted for the largest proportion, followed by signaling and cellular processes. Comparing the two groups in the subcategory of KEGG function, disease-associated genes had a larger proportion involved in amino acid metabolism, antimicrobial resistance and transcription factors, while carbohydrate metabolism had a high ratio of health-related genes.

Among the differentially expressed genes between strains from diseased and healthy sources, we found some key disease-related genes, including the virulence genes *essD*, *uhpt*, *sdrF*, *sdrG*, *fbe*, and *icaABCDR* and the transcriptional regulators *lrpC* and *cysL* (**Figure 5A**). *SdrF*, *sdrG* and *fbe,* which are microbial surface components recognizing adhesive matrix molecules (MSCRAMMs), showed homology with the virulence gene

**FIGURE 5.**
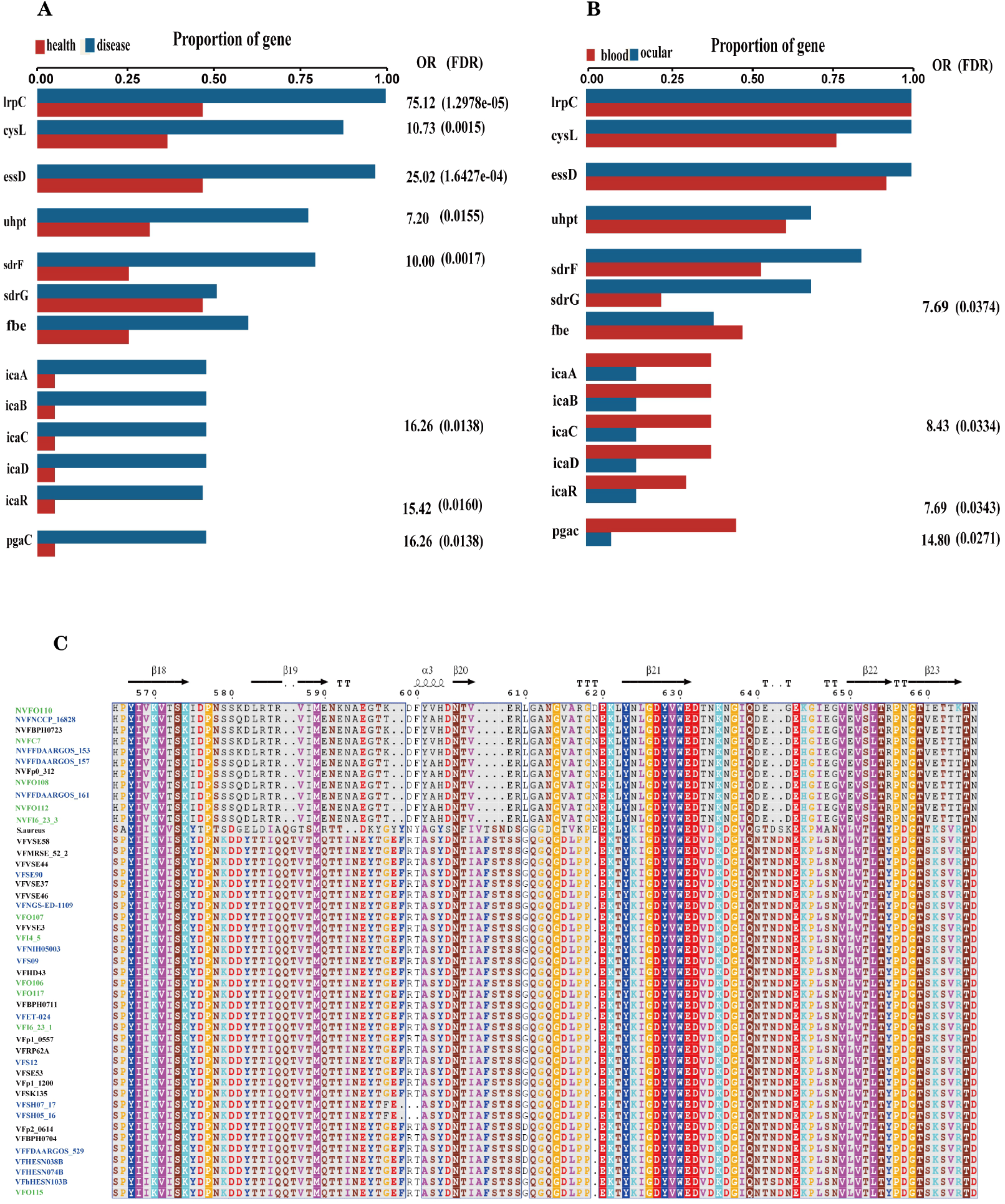
Key genes associated with diseased source pathogens. A. Frequency of key genes among bacteria isolated from different health state. OR, odds ratio for association between the presence of the genes and disease versus health source; P values were calculated using Fisher’s exact test. B. Frequency of key genes among bacteria isolated from different human source. OR, odds ratio for association between the presence of the genes and disease versus health source; P values were calculated using Fisher’s exact test. C. Multiple alignment of virulence gene sdrG and non virulence gene sdrG protein sequences across the collected blood source S.epidermidis (blue), intraocular isolates (green), staphylococcus aureus and sdrG positive skin source strains.

*sdrE*, but not all of the gene sequences were identified as virulence factors. By multiple alignment with Clustal omega, we found that compared to both the virulence gene *sdrG* protein sequences and *Staphylococcus aureus*, nonvirulence gene *sdrG* protein sequences mutated in the β-strand (**Figure 5C**), which may decrease virulence.

Interestingly, to compare genes between ocular and other places of organism, we found that biofilm formation-related genes *icaABCDR* and *pgaC* were markedly enriched in blood but not in ocular tissue (**Figure 5B**). Moreover, although *sdrG* seemingly enriched in ocular cells, the β-strand mutation rate reached 42.86% (6/14), which may lead to weaker adherence ability of the microbes to components of the extracellular matrix of the host, suggesting that biofilm formation may not be a direct factor for intraocular infection of *S. epidermidis*.

### Antimicrobial Resistance across *S. epidermidis*

Antimicrobial resistance is very common among *S. epidermidis* isolates, and genes within *S. epidermidis* involved in antibiotic resistance contribute to the persistence of clinical infection and often limit treatment options(71). To investigate antimicrobial resistance across *S. epidermidis*, we analyzed all known AMR genes within our 187 genomic datasets. According to our analysis of the ResFinder and CARD databases, we found 41 different genes involved in resistance to 19 antibiotics **(Figure 6)**. Nearly all isolates carried at least three antibiotic resistance genes. Among the genes involved in AMR, our data showed that two genes, *norA*, which is associated with fluoroquinolone antibiotics, from the AMR gene family with major facilitator superfamily (MFS) antibiotic efflux pumps and *dfrC*, a diaminopyrimidine antibiotic-associated gene, were conserved in all strains. This result was consistent with the data in the comprehensive antibiotic resistance database (CADR: https://card.mcmaster.ca/ontology).

**FIGURE 6.**
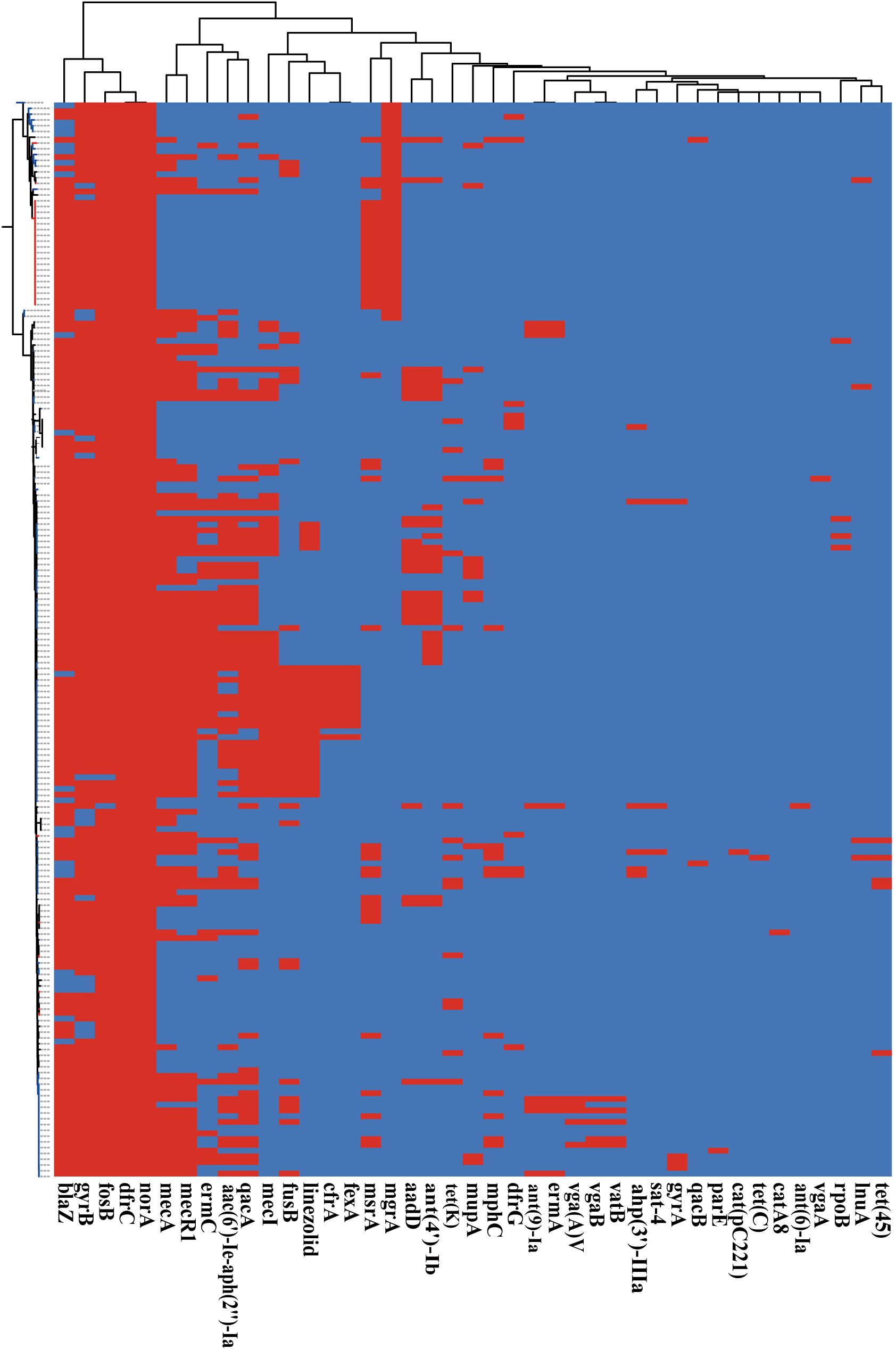
Presence of antibiotic resistant genes across Staphylococcus epidermidis strains. Heatmap indicates of 41 antibiotic resistant genes involved. The red color stands for genes that existed and the dark blue color for missing ones.

Based on Fisher’s exact test of strains from different host health states and niches, we found that the methicillin resistance genes *mecA* and *mecR1* (FDR <0.01), fluoroquinolone antibiotic gene *ace* (FDR <0.05) and aminoglycoside antibiotic gene *aac(6’)-Ie-aph(2’’)-Ia* (FDR <0.001) were enriched in strains isolated from diseased hosts. The presence of *mecA* was consistent with the SCC*mec*-positive strains because the mobile genetic element SCC*mec* carried the gene *mecA*. Antibiotic susceptibility testing currently uses oxacillin and cefoxitin to assess resistance to methicillin, and the results showed that the intraocular strain oxacillin resistance rate was 5/11 (**Table 2**), which was consistent with the results for cefoxitin and *mecA* gene carriers. Moreover, strains from healthy skin had significant enrichment of the msrA and mgrA genes with multidrug resistance (FDR <0.001), while isolates from the ocular, blood and respiratory tracts had no significantly enriched antibiotic resistance genes, which suggested that regardless of niche sources, none of these isolates were resistant to specific antibiotics at the genetic level.

**Table2.**
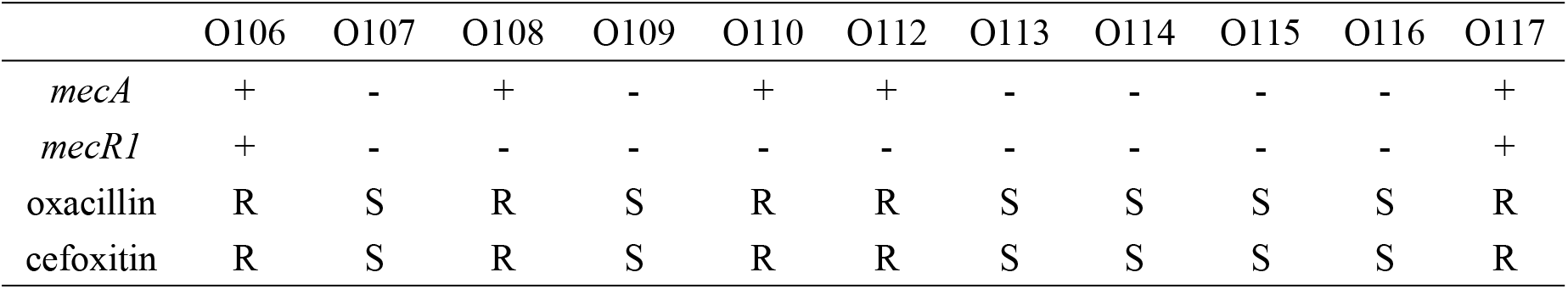
*Mec* gene carried rate and antibiotic susceptibility testing results of 11 ocular trauma-source *Staphylococcus epidermidis*

## Discussion

Coagulase-negative *Staphylococcus* was the most common pathogen in traumatic endophthalmitis(72). *S. epidermidis*, which colonizes the normal mucosa, skin flora and intraocular tissue of humans and other mammals, is the most common coagulase-negative gram-positive *Staphylococcus*(69)and one of the major leading causes of clinical infections. What still needs to be discussed is whether *S. epidermidis* is isolated as a pathogen or contaminant. Although the genome characteristics of *S. epidermidis* have been studied in recent years, a comprehensive understanding of the genomes of ocular-sourced strains is lacking. To the best of our knowledge, this is the first and largest collection of bacterial genome sequences isolated from patients with *S. epidermidis* intraocularly. By collecting, sequencing and analyzing the genomes of eleven intraocular isolates and incorporating publicly available genomes for *S. epidermidis*, we gained insight into the phylogenetic and molecular characteristics of intraocular and other niche pathogens. The genetic differences between pathogenic and commensal *S. epidermidis* were also investigated comprehensively.

In this study, we compared the phylogenetic diversity and genome characteristics of *S. epidermidis* from different niches and different host health states. The host niches, including the eye, blood, skin and respiratory tract, were all from diseased hosts. The 11 ocular-sourced genomes we sequenced revealed similar high-quality benchmark data, including genome size, GC content, and the number of predicted genes, which were similar to the data we collected from public databases and had a relatively compact genome with an average size of approximately 2.58 Mb. Furthermore, the completeness of each whole-genome sequencing exceeded 99%, and the contamination level of the sequencing library was also very low. The high coverage of genome assembly helped us obtain a complete and accurate genome. Consistent with the results of previous studies (27)(39)(41), whether in the pangenome analysis of all 187 strains or strains of various niches, including ocular sources, all showed that the size of the *S. epidermidis* genome was relatively constant, extracted from a larger gene pool, indicating an increase in the “open” pangenome; with each genome sequence, some new genes were added. To some extent, the pangenomic state (“open” or “closed”) of an organism depends in part on its ability to obtain exogenous DNA(64); for example, the large number of genes involved in the mobile group makes horizontal gene transfer between staphylococcal stains easier, and mobile genetic elements such as SCC*mec*, ACME, and plasmids lead to an increase in the “open” pangenome(73). The phylogenetic tree constructed based on genome-wide core SNPs reveals important details that were not available in the use of traditional single gene markers (16S rDNA) or multilocus sequence typing (MLST)(40). From the analysis of the phylogenetic tree, we found that 187 strains of *S. epidermidis* formed two distinct clusters with different pathogenic abilities, and clinically pathogenic strains were generally ST5 and ST2, while the ST691 strains were derived from healthy skin. The earlier reported phenomenon of evolutionary distance between ST2 *S. epidermidis* was extremely short(40) and was also observed in the ST5 and ST691 strains.

It is well established that *S. epidermidis* is a common human skin commensal; if it breaks through the surface of the skin and enters the blood, it is considered a pathogen(12). Similarly, *S. epidermidis* is an important commensal on the ocular surface(74), but usually caused by trauma, it can enter the eyes to cause intraocular infection. Through whole-genome analysis, we have provided a powerful framework to redefine species clustering in the genus, locate genetic traits, and rate the importance of disease-causing genes based on their presence or absence. Using comparative genomics methods to analyze the genomes of *S. epidermidis* from different sources, we obtained the potential pathogenic marker genes *lrpC, cysL, essD, uhpt, sdrF, sdrG, fbe, and icaABCDR*. *LrpC* and *cysL* both have helix-turn-helix (HTH)-type transcriptional regulators. The *lrp*-like regulatory factor consisted of a helix-turn-helix (HTH)-type n-terminal DNA binding domain, which was connected to the C-terminal RAM domain (amino acid metabolism regulation) and was responsible for cofactor binding and oligomerization(75). Thus, *lrpC* was a transcriptional regulator with a possible role in the regulation of amino acid metabolism and the growth phase transition. CysL belongs to the *lysR* family transcriptional regulator. The *lysR* HTH domain is a DNA-binding, winged helix-turn-helix (wHTH) domain consisting of approximately 60 residues in the *lysR*-type transcription regulator (LTTR). LTTR is one of the most common regulatory factor families in prokaryotes. The c-terminus of the *lysR* protein can contain a regulatory domain with two subdomains, which participate in (1) coinducer recognition/reaction and (2) DNA binding and response(76). LTTRs can activate the transcription of operons and regulons involved in the regulation of various functions, such as amino acid biosynthesis, CO2 fixation, antibiotic resistance, virulence factor regulation, nitrogen-fixing bacterial nodulation, oxidative stress response or aromatic compound catabolism. However, the specific functions of the transcriptional regulators *cysL* and *lrpC* in *S. epidermidis* need to be further studied.

In this work, pathogenic marker genes *essD*, *uhpt*, *sdrF*, *sdrG*, *fbe*, and *icaABCDR* were identified as virulence genes. Interestingly, we found that the potential virulence genes *essD* and uhpt had high homology with *esaD* and *hpt* in *Staphylococcus aureus*. *EsaD* is a type VII secretion system secreted protein, a nuclease toxin, which may play a key role in bacterial competition(77). Hexose phosphate is an important carbon source in the cytoplasm of host cells(78). Bacterial pathogens invade, survive and breed in different host epithelial cells, utilizing hexose phosphate in the host cytoplasm to obtain energy and synthesize cell components through the hexose phosphate transport (HPT) system (78). The HPT system of *S. aureus*, which includes the *hptRS* (a new type of two-component regulatory system), *hptA* (a phosphate sensor) and *uhpT* (a hexose phosphate transporter) genes, made it stable in the host cell and may be an important target for the development of new anti-*staphylococcal* therapies(78). The potential virulence genes essD and uhpt of *S. epidermidis* identified in this study could be due to horizontal gene transfer between *Staphylococcus* strains; for example, the genomes of *S. epidermidis* and *S. aureus* exchanged during evolution. Furthermore, the genes *sdrF, sdrG, fbe*, and polysaccharide intercellular adhesion gene (*icaABCD*) were considered to be related to biofilm formation(28). The formation of biofilms was the main virulence factor *S. epidermidis* contributed to the persistence of clinical infections(23). Here, the seven genes all pertain to adhesive molecules, which are well-known factors involved in biofilm formation. *SdrF, sdrG* and *fbe* are a subset of microbial surface components recognizing adhesive matrix molecules (MSCRAMMs)(28); they were covalently anchored to the cell wall and characterized by a segment composed of repeated serine aspartate (SD) dipeptides(79). MSCRAMMs are bacterial surface proteins that mediate the adhesion of microorganisms to the host’s extracellular matrix components(79). By sequence analysis, we found that not all of the *sdr* gene sequences were identified as virulence factors. Through further multiple sequence alignment, we found that the virulence sequence of *sdrF* was very different from the nonvirulence sequence, including the length of the sequence, while only the β chain mutation occurred between the two *sdrG* genes. We suspected that the occurrence of these mutations may reduce the virulence of *sdrG*, and it was clear that mutagenesis is overrepresented in intraocular isolates compared to blood isolates. According to our enrichment analysis, we found it was possible to differentiate the intralocular pathogenic strains from those of blood, with biomarkers related to biofilm formation polysaccharide intercellular adhesion *icaABCD* enriched in blood but not in ocular tissue. This may be related to the fact that *S. epidermidis* was more likely to form biofilms on medical devices such as catheters and artificial heart valves(2). Bacterial cells on these devices can break away from the biofilm and enter the bloodstream, leading to bacteremia, increasing morbidity and potential mortality(2). On the other hand, this also reflected that the intraocular infection of *S. epidermidis* was not directly related to its biofilm formation. Of course, a larger amount of data and further experiments are required to prove this point.

*S. epidermidis* has also been found to be a treasure trove of antibiotic resistance(27). Through a rare horizontal gene transfer event, its determinants of toxicity were shared with other more pathogenic species, such as *S. aureus*, as confirmed in previous studies(80). In our study, we found that 94/187 (50.2%) of the collected *S. epidermidis* species were methicillin-resistant *Staphylococcus epidermidis* (MRSE) with the *mecA* gene and SCC*mec* elements. In particular, the SCC*mec* cassette that conferred β-lactam resistance was often transferred between staphylococcal strains, enabling them to rapidly evolve and adapt to antibiotic selection pressures and providing additional competitive advantages. This provided strong support for the idea that pathogens influence the risk of infection by the background microbiota through HGT of the pathogenic host, thereby increasing the risk of infection in other parts of the body, such as methicillin-resistant *S. aureus* in the nasal cavity affecting *S. epidermidis*-infected endophthalmitis(40).

## Conclusion

Our study provided information on the molecular characteristics of different pathogenic *Staphylococcus epidermidis* isolated from different host niches around the world, including ocular strains that were overlooked in previous studies. Pangenome and phylogenetic analysis that *S. epidermidis* had an open pangenome and two founder lineages that had different pathogenicities. Interestingly, methicillin-resistant *S. epidermidis* was concentrated in the pathogenic branch. Although the endophthalmitis-associated *S. epidermidis* isolated in this study was relatively dispersed in evolution, they were close to the clinical pathogenic strains, which proved their pathogenicity. Based on comparative genomics, we identified 8 potential biomarkers related to strain pathogenicity and provided evidence that horizontal gene transfer (HGT) may occur between *Staphylococcus* strains. Moreover, we reported the complete genome sequence of *S. epidermidis* that caused traumatic endophthalmitis and found that those strains causing intraocular infection may be independent of biofilm formation. Overall, this study revealed genetic diversity and pathogenic differences in different sources of *S. epidermidis*.

